# Using muscle-tendon load limits to assess unphysiological musculoskeletal model deformation and Hill-type muscle parameter choice

**DOI:** 10.1101/2024.04.18.590034

**Authors:** Lennart V. Nölle, Isabell Wochner, Maria Hammer, Syn Schmitt

## Abstract

Musculoskeletal simulations are a useful tool for improving our understanding of the human body. However, the physiological validity of predicted kinematics and forces is highly dependent upon the correct calibration of muscle parameters and the structural integrity of a model’s internal skeletal structure. In this study, we show how ill-tuned muscle parameters and unphysiological deformations of a model’s skeletal structure can be detected by using muscle elements as sensors with which modelling and parameterization inconsistencies can be identified through muscle and tendon strain injury assessment.

To illustrate our approach, two modelling issues were recreated. First, a model repositioning simulation using the THUMS AM50 occupant model version 5.03 was performed to show how internal model deformations can occur during a change of model posture. Second, the muscle material parameters of the OpenSim gait2354 model were varied to illustrate how unphysiological muscle forces can arise if material parameters are inadequately calibrated. The simulations were assessed for muscle and tendon strain injuries using previously published injury criteria and a newly developed method to determine tendon strain injury threshold values.

Muscle strain injuries in the left and right *musculus pronator teres* were detected during the model repositioning. This straining was caused by an unphysiologically large gap (12.92 mm) that had formed in the elbow joint. Similarly, muscle and tendon strain injuries were detected in the modified right-hand *musculus gastrocnemius medialis* of the gait2354 model where an unphysiological reduction of the tendon slack length introduced large pre-strain of the muscle-tendon-unit.

The results of this work show that the proposed method can quantify the internal distortion behaviour of musculoskeletal human body models and the validity of Hill-type muscle parameter choice via strain injury assessment. Furthermore, we highlight possible actions to avoid the presented issues and inconsistencies in literature data concerning the material characteristics of human tendons.

## 1 Introduction

Musculoskeletal simulations are a useful tool for improving our understanding of movement generation [1] and the body’s response to external loads [2]. However, while the underlying mathematical methods of such simulations can be verified in the sense that equations are solved correctly, establishing their physiological validity has proven to be a challenging task [3,4]. A defining trait of musculoskeletal models is the inclusion of active muscle elements in the form of so-called Hill-type muscles [5], which are used to generate movements during the simulation runtime. Many different variants of Hill-type muscles exist [6], each designed to improve on the original material model. However, despite the constant progress in the development of Hill-type muscle models, they remain sensitive to parameter variations [7–10] and can be prone to numerical instability [11]. In the context of finite element (FE) models, further challenges are presented by the models’ inherent deformability, which can cause joints to shift or dislocate during repositioning simulations required to adjust the model’s posture [12–14]. As a result, the physiological validity of the predicted kinematic responses and internal forces is highly dependent upon the correct calibration of muscle parameters and, in the case of FE models, the structural integrity of their internal skeletal structure. But how can unphysiological muscle parameter sets or model deformations be identified? In our current study, we try to answer this question by using injury criteria to detect ill-tuned muscle parameters and deformations of a model’s skeletal structure which may lead to unrealistic model behaviour. This approach is based on the stipulation that during physiological movements, no injury of any kind should occur in the musculoskeletal model.

In the past, numerous injury criteria and injury threshold values for hard tissue injuries of the skull [15], nose [16–18], ribs [19], pelvis [20,21] and leg [22–26] as well as soft tissue injuries of the brain [27–31], neck ligaments [18,32,33], intervertebral discs [34], lungs [35], heart [36], kidney [37], spleen [38], liver [37,38], intestines [36], blood vessels [36] and ankle ligaments [39,40] have been proposed. While these injury criteria have proven useful especially in the field of automotive safety [41], they cannot be applied to all musculoskeletal models. For example, multibody (MB) models lack deformable structures, necessary for the evaluation of many soft and hard tissue injuries. In our previous studies, two novel injury criteria for assessing the severity of muscle and tendon strain injuries, called the Muscle Strain Injury Criterion (MSIC) [42] and the Tendon Strain Injury Criterion (TSIC) [43], were developed. These criteria allow for the interpretation and evaluation of forces and strains occurring in the muscle-tendon-unit (MTU) and, as such, can be applied to any musculoskeletal model which includes a muscle material model capable of predicting MTU forces and tendon strains. Because of their general applicability, we will show how unphysiological musculoskeletal model deformation and Hill-type muscle parameter choice can be assessed using the MSIC and TSIC. Through applying this knowledge on the load limits of the MTU, each muscle element can thus act as a sensor with which modelling and parameterization inconsistencies can be identified.

To illustrate our proposed method, the previously outlined modelling issues were recreated using two well-established musculoskeletal models in typical load cases. First, an exemplary FE model repositioning simulation using the Total Human Model for Safety (THUMS) AM50 occupant model version 5.03 [44] was performed in LS-DYNA (Ansys, Canonsburg, PA, United States), to show how internal model deformations can be detected. Second, the muscle parameters of the OpenSim [45] gait2354 model [46–49] were varied to demonstrate the efficacy of our method in identifying unphysiological parameter sets. Finally, possible actions to avoid the presented issues, as well as inconsistencies in literature data concerning the material characteristics of human tendons, are highlighted.

## 2 Materials and Methods

### 2.1 Description of models and load cases

#### 2.1.1 Repositioning FE simulation

The FE repositioning simulation was performed using the THUMS AM50 occupant model version5.03 [44]. Seat, steering wheel and pedals were taken from the openly available THUMS version 5.03 validation catalogue example depicting a whole body frontal sled test [50,51]. The muscle material of relevant muscles spanning the elbow joint was changed from the LS-DYNA internal Hill-type material *MAT_MUSCLE to a more physiological Hill-type muscle variant developed by Günther et al. [52] and Häufle et al. [53], which aims to more realistically represent the eccentric force–velocity relation of the biological muscle. Additionally, it consists of distinct muscle and tendon sections, which can be individually evaluated for injury with the MSIC and the TSIC. This Hill-type muscle material was initially implemented in LS-DYNA as the so-called extended Hill-type muscle (EHTM) material model by Kleinbach et al. [54] and has since been updated by Wochner et al. [55] and Martynenko et al. [56]. In our current study, the functionality of the EHTM was further expanded by adding an inbuilt strain injury assessment functionality using the MSIC and TSIC. This version 4.0.00 of the EHTM has been made publicly available in a data repository maintained by Nölle et al. [57]. In both arms, *musculus brachialis, musculus biceps brachii* (long and short head), *musculus brachioradialis, musculus triceps brachii* (long, lateral, and medial head), *musculus anconeus* and *musculus pronator teres* were replaced, resulting in 18 total material changes in the THUMS model. The original *MAT_MUSCLE material parameters were converted for the use with the EHTM using the methodology described in Wochner et al. [55] and Nölle et al. [57]. A list of all edited muscles and their material parameters is given in S-Table 1 and S-Table 2 in S1 File.

**Table 1.** Minor TSIC thresholds of positional, TMM, EHTM and energy storage tendons.

| Tendon type | Positional tendon | TMM | EHTM | Energy storage tendon |
| --- | --- | --- | --- | --- |
| Minor TSIC threshold [% strain] | 2.8 | 5.16 | 9.35 | 14.4 |
| $F/F_{\max}$ at threshold | 2.17 | 1.97 | 1.60 | 1.16 |
| Slope [1/%] | 1.33 | 0.52 | 0.24 | 0.10 |
| y-intercept | -1.53 | -0.71 | -0.60 | -0.30 |

The default THUMS model posture (Fig 1A) was altered through a repositioning simulation which is described in the following and aimed to achieve an extended posture of the arms needed to adapt the model for the use in an alternative interior configuration (Fig 1B). The arm extension was kept within a physiological range of motion, as to not purposefully overextend the arms. For the repositioning, the hands were constrained to the steering wheel using a tied contact, while the model’s torso was fixed in space. The steering wheel was then moved in four steps. First, the wheel was displaced 100 mm upwards in positive z-coordinate direction from t = 0.0 s to t = 0.4 s with a constant speed of 250 mm/s. Second, the model was allowed to settle for 0.1 s from t = 0.4 s to t = 0.5 s. Third, the steering wheel was moved 150 mm forward in positive x-direction from t = 0.5 s to t = 1.1 s with a constant speed of 250 mm/s. This was followed by a final settling period from t = 1.1 s to t = 1.2 s.

**Fig 1.**
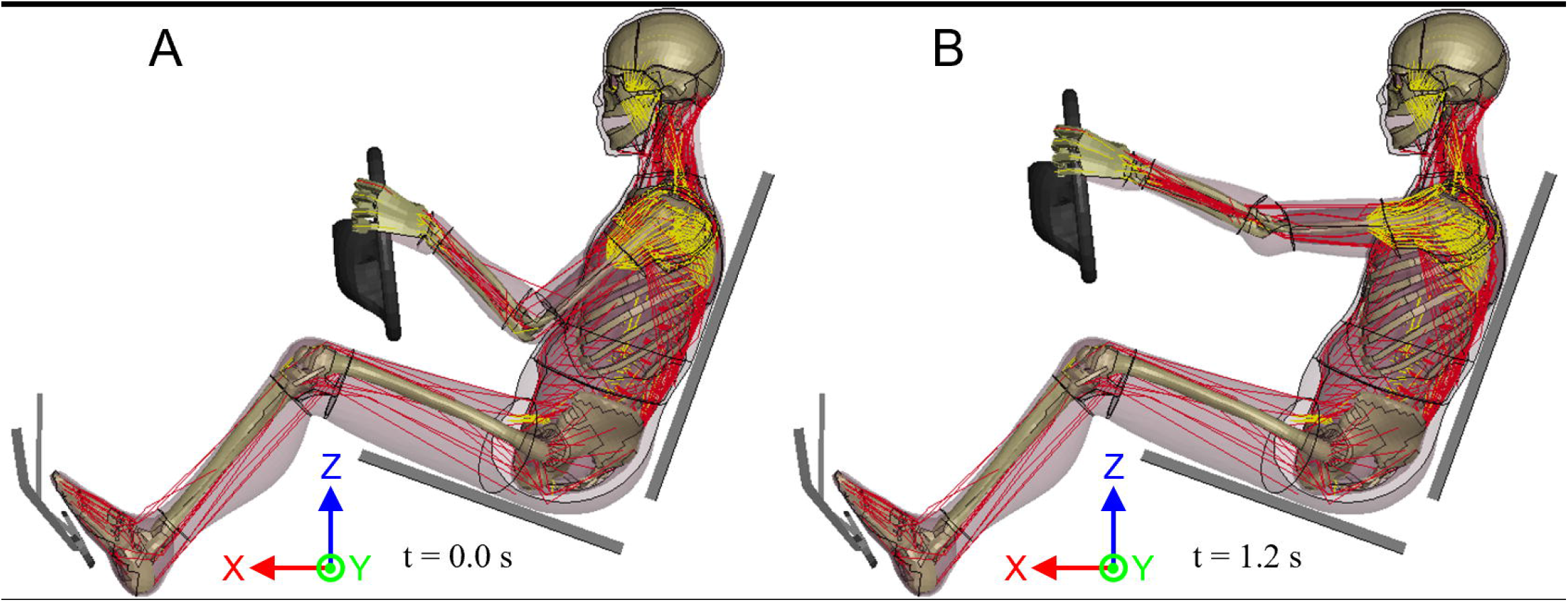
Visualisation of the THUMS version 5.03 occupant model. (A) Initial THUMS occupant position at t = 0.0 s ; (B) Repositioned THUMS with extended arms at t = 1.2 s.

The repositioning simulation was performed with a user-compiled double precision (DP) symmetric multiprocessing (SMP) version of LS-DYNA R9.3.1 (Ansys, Canonsburg, PA, United States) including EHTM version 4.0.00 [57]. The simulations were run on a high-performance workstation equipped with an AMD Ryzen Threadripper 3990X 64-core processor (AMD, Santa Clara, CA, United States) using 32 SMP threads. All visualisations of the FE simulation were done using LS-PrePost V4.8.24 (Ansys, Canonsburg, PA, United States). Only the 18 changed EHTM muscles were assessed for muscle and tendon strain injury occurrence.

#### 2.1.2 Gait cycle MB simulation

To illustrate the detection of ill-tuned muscles in an MB context, two sets of partial gait cycle simulations were performed in OpenSim [45] using the gait2354 model [46–49]. For the first set, necessary muscle control parameters were determined with the OpenSim Computed Muscle Control (CMC) functionality [58] following the steps from the OpenSim online documentation [59]. The resulting CMC outputs where then used in a forward dynamics simulation, following the protocol similarly listed online [60], to output the muscle analysis data needed for the injury assessment. The partial gait cycle started at t = 0.83 s with the toe-off of the left foot and ended shortly before the left heel-strike at t = 1.08 s as pre-defined in the CMC and forward dynamics simulation setups. For the second set of simulations, this exact process was repeated with one modification, namely a change in the parameterisation of a singular muscle to purposefully introduce an exemplary modelling error. As such, the tendon slack length of the right *musculus gastrocnemius medialis* head was reduced by 20% to 0.288 m from its original length at 0.361 m. The resulting partial gait cycles of both simulation sets are displayed in Fig 2. All simulations were run in OpenSim version 4.4 [45,61] on an Intel® Core™ i7-10510U processor. Since the gait2354 model uses the inbuilt Thelen 2003 muscle model (TMM) [62], in which tendon and muscle belly are considered separately like in the EHTM, all muscles present in the default and modified gait2354 models were evaluated for strain injuries.

**Fig 2.**
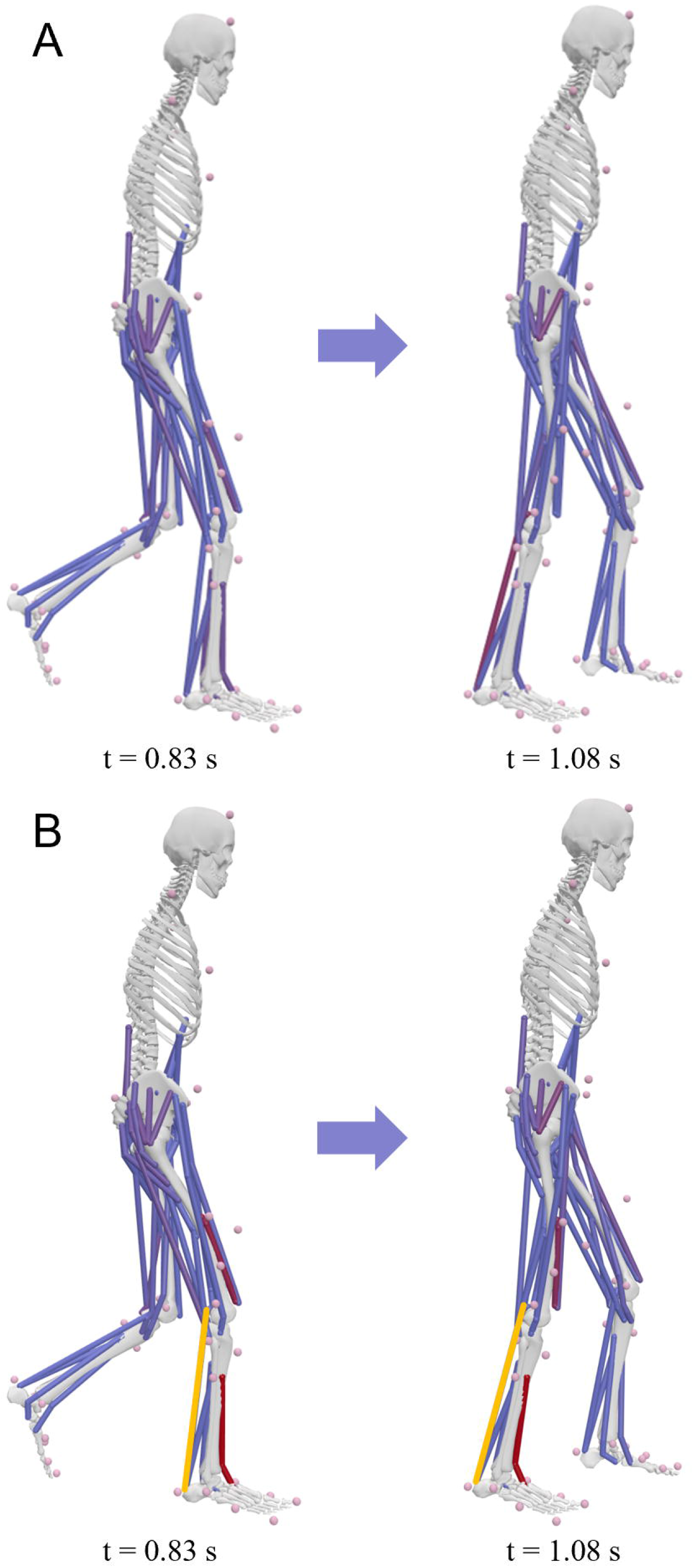
Forward dynamics gait cycles of the gait2354 models. (A) the default gait2354 model; (B) the modified gait2354 model. The differently parameterised *musculus gastrocnemius medialis* is highlighted in yellow.

### 2.2 Applied injury criteria

The assessment of muscle and tendon strain injury severity was performed using two injury criteria, the MSIC [42] and the TSIC [43]. Both criteria categorise the grade of the sustained strain injury based on the deformation stages of the muscle and tendon (Fig 3) as previous studies [63–66] have shown that the regions of the stress-strain curve can be linked to the degree of tissue damage. As such, the injury criteria encompass three strain injury thresholds, with the minor injury threshold set at the start of the strain hardening region, the major injury threshold at the start of the necking region, and the rupture threshold at the point of material failure. For this study, only the minor injury thresholds of MSIC and TSIC were used to evaluate the simulation results, as movements within physiological ranges of motion should not result in any form of strain injury regardless of severity. Additionally, the tendons of the TMM and the EHTM only reproduce the non-linear toe region and an infinitely continued linear elastic region of the tendon stress-strain curve, which means that in a strict sense, both muscle material models lose their validity for large strains at which plastic deformation is expected, and where major strain injuries would occur.

**Fig 3.**
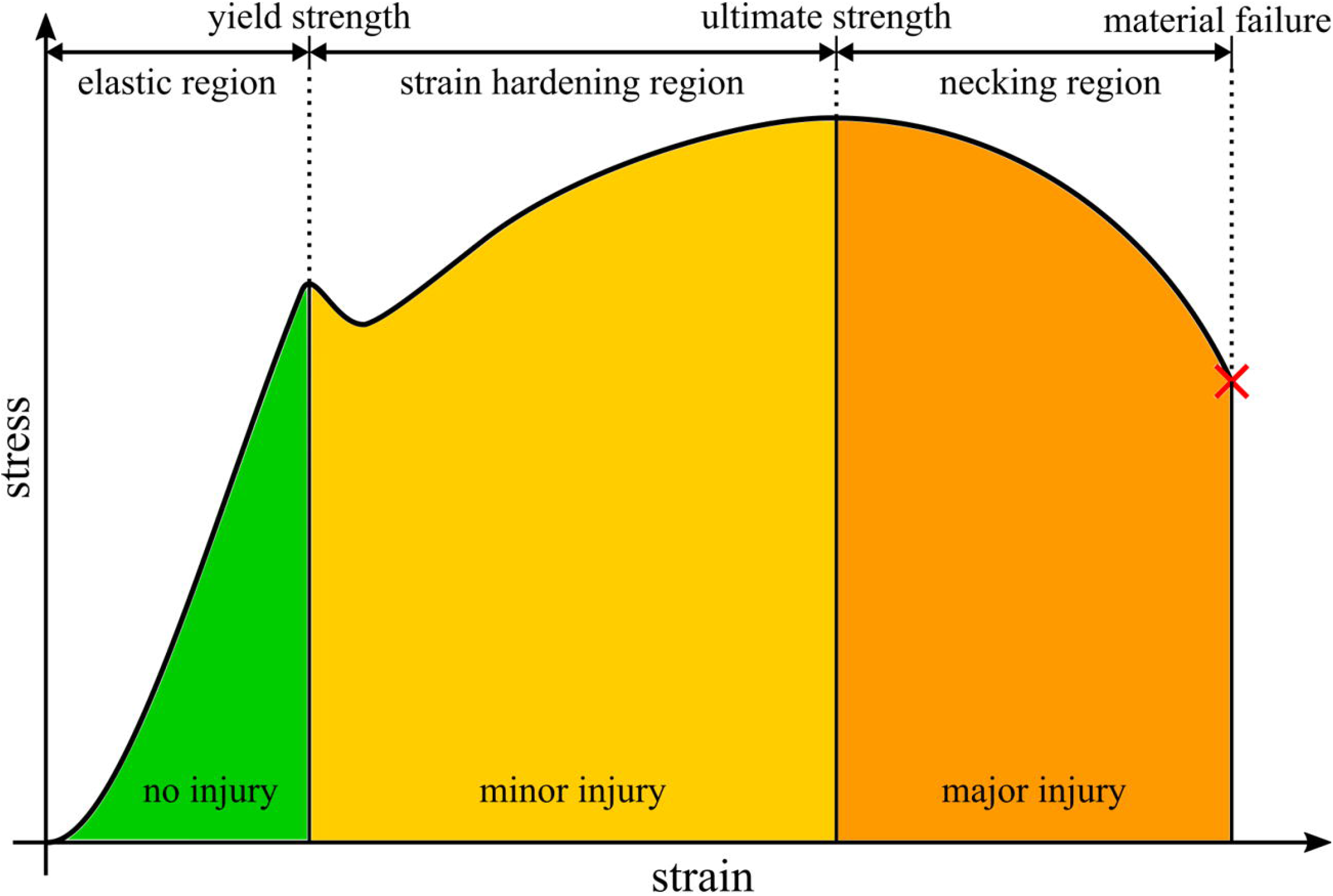
Schematic stress-strain curve with highlighted injury thresholds.

The reliability of the injury assessment via both MSIC and TSIC strongly depends on the correct choice of MTU material parameters. The force based MSIC thresholds scale with each muscle’s maximum isometric force (F_max_) and the muscular activity (a) while muscle rupture is defined to occur at them force to failure F_tf_ = 3 F_max_ [42]. Therefore, they can be easily applied to any muscle, with their inherent scalability eliminating the chance of a mismatch between the muscle material parameters and the defined injury thresholds. In contrast, only one fixed set of strain based TSIC injury thresholds has been proposed thus far [43]. This limits the applicability of the criterion to a subset of tendons which exactly match the deformation characteristics of the stress-strain curve used to define the TSIC threshold values. For example, highly compliant tendons might reach their plastic deformation stage at strains that would have already ruptured stiffer tendons. As such, the injury severity of tendon strains cannot be reliably evaluated with constant, generic thresholds.

In an attempt to provide a simple functional classification of tendon types, Kaya et al. [67] distinguish two types of tendons. The so called ‘energy storage tendons’ serve to store and release kinetic energy during dynamic movements and thus need to be more elastic than their stiffer counterpart, the ‘positional tendon’, whose main function is to ensure a near lossless transfer of forces between muscle and bone. However, not all tendons fit these two archetypical tendon types perfectly, resulting in a multitude of sensible tendon parameterisations within a corridor between maximally stiff or maximally compliant. Therefore, this work introduces a method to determine minor TSIC thresholds for tendons with infinitely linear deformation characteristics. For this, two boundary curves with known yield strength points, at which the material deformation switches from elastic to plastic, are defined. These boundary curves represent an ideal positional or energy storage tendon, respectively. The yield strength points of the boundary curves then serve as the grid points of a line linearly interpolated according to Eq 1.

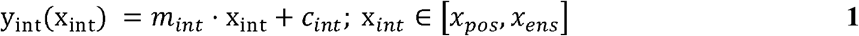

with

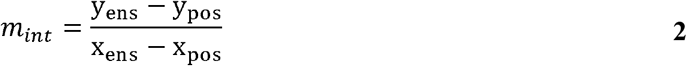

and

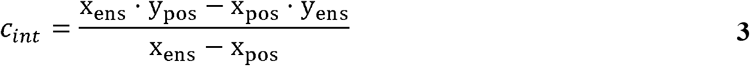

where x_int_ and y_int_ are the abscissa and ordinate values of the interpolated line, m_int_ is the slope of the interpolated line, c_int_ is the y-intercept of the interpolated line, x_ens_ and y_ens_ are the abscissa and ordinate values of the energy storage tendon yield strength point and x_pos_ and y_pos_ are the abscissa and ordinate values of the positional tendon yield strength point.

The yield strength points of tendons can then be determined by finding the intersection between the linear part of the tendon deformation curve and the interpolated yield strength line (Eq 4).

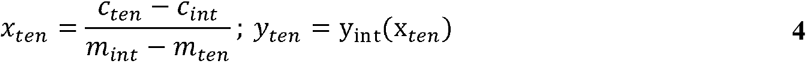

where x_ten_ and y_ten_ are the abscissa and ordinate values of the tendon’s yield strength point, m_ten_ is the slope of the tendon curve and c_ten_ is the y-intercept of the tendon curve.

To set the boundary curves used in our current work, data on what could be considered ideal energy storage and positional tendons was collected from literature in a first step. Experimental data on the deformation characteristics of human energy storage tendons can be found in the work of Shaw and Lewis [68], who studied the tensile properties of the human Achilles tendon and determined a suitable constitutive relation between its strain and the resulting stress (Eq 5).

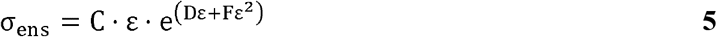

where σ _ens_ is the stress of the energy storage Achilles tendon, ε is the strain in percent and C, D and F are material constants.

For this work, the constant values were set to match those reported for the donor age group from 36 to 50 years old as C = 1.386 MPa, D = 0.123 and F = -0.004 [68]. The Achilles tendon was deemed to be a suitable representative for the energy storage tendon category, as this tendon is among the most elastically deformable in the human body with reported Young’s moduli between E = 459 MPa [68] and E = 822 MPa [69]. Similarly, tensile test data derived from unenbalmed human foot flexor tendons [70] were used to represent the deformation characteristics of positional tendons, as they are exceedingly stiff (E = 2.4 GPa [70]) to translate the contraction of relatively short muscles along a comparatively long tendon to create toe movement. The stress-strain curves of the Achilles tendon (energy storage tendon) calculated using Eq 5 and the average foot flexor tendon taken from Benedict et al. [70] as the positional tendon stress σ_pos_ are shown in (S-Fig 1 in S1 File).

Next, the energy storage and positional stress-strain curves were converted to force-strain curves as the tendons of the EHTM and TMM models do not output stresses but forces instead. this stress to force conversion was performed according to Eq 6.

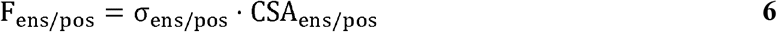

where F_ens/pos_ is the force of the energy storage or positional tendon,σ _ens/pos_ is the stress of the energy storage or positional tendon, and CSA_ens/pos_ is the cross-sectional area of the energy storage or positional tendon.

Since the necessary data on the cross-sectional area (CSA) of the Achilles and foot flexor tendons needed for this transfer was not provided in the original publications, they were instead taken from literature as CSA_ens_ = 60.90 mm^2^ [71] and CSA_pos_= 10.32 mm^2^ [72]. The derived force-strain curves are depicted in S-Fig 2in S1 File.

Following the stress-to-force conversion, the force-strain curves were normalised to the maximum isometric forces acting on the tendons (Eq 7).

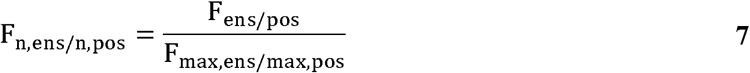

where F_n,ens/n,pos_ is the normalised force of the energy storage or positional tendon, F_ens/pos_ is the force of the energy storage or positional tendon, and F_max,ens/max,pos_ is the maximum isometric force associated with the energy storage or positional tendon.

For the positional foot flexor tendons, an average F_max,pos_ of 315 N was calculated based on data from Saraswat et al. [73], while the sum of forces acting on the energy storage Achilles tendon was defined as F_max,ens_ = 5500.3 N by adding up the F_max_ values of *musculus gastrocnemius* (*lateralis* and *medialis*) and *musculus soleus* reported by Arnold et al. [74]. However, normalising the force-length curves to these F_max_ values yielded results (S-Fig 3 in S1 File) which are either incongruous with other experimental works or physiologically unrealistic. The positional tendon ultimate strength of 2.05 times F_max_ appears to be uncharacteristically low, given that fact that the experiments from Hasselman et al. [75] and Noonan et al. [76] show that mammalian tendons can transmit forces exceeding 3 times F_max_. Similarly, the normalised energy storage tendon reaches its ultimate strength at just 0.73 times F_max_, indicating that the Achilles tendon could not even withstand the maximum isometric forces of the connected muscles. Consequently, any somewhat athletic movement involving the Achilles tendon should cause it to rupture instantly.

The authors therefore decided to scale the positional and energy storage force-strain curves using scale factors c_ens/pos_ calculated according to Eq 8.

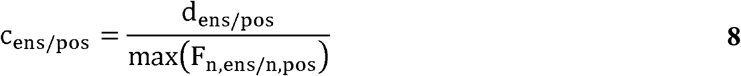

where c_ens/pos_ is the scale factor of energy storage or positional tendons, d_ens/pos_ is the desired proportionality factor with regards to F_max_, and max(F_n,ens/n,pos_) is the maximum value of the normalised energy storage or positional tendon force.

This resulted in c_ens_ = 2.05 for d_ens_ = 1.5 and c_pos_ = 1.47 for d_pos_ = 3 to achieve ultimate strength values of 1.5 times F_max_ and 3 times F_max_ respectively. The force curves where then scaled using Eq 9.

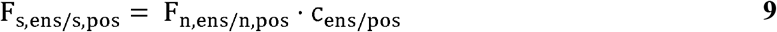

where F_s,ens/s,pos_ is the scaled, normalised force of the energy storage or positional tendon, F_n,ens/n,pos_ is the normalised force of the energy storage or positional tendon, and c_ens/pos_ is the scale factor of energy storage or positional tendons.

This ensured that the general quality of the curves and their comparative magnitude was preserved while increasing the injury resistance of the energy storage tendon to account for forces above F_max_ as they appear during eccentric contractions [77]. Additionally, the redefined positional tendons allow the transmission of forces in accordance with the previous findings by Hasselman et al. [75] and Noonan et al. [76] on which the MSIC was based. Ensuring this compatibility with the MSIC thresholds is vital, as muscle rupture is defined to occur at the force to failure F_tf_ = 3 F_max_ [42] and without the scaling of the force-strain curves, these forces could never occur in the muscle but would instead always cause tendon rupture first.

After the force scaling, the minor TSIC thresholds for the EHTM and TMM tendon were determined in a final step. For this, the minor injury thresholds of the positional and energy storage tendons were found by determining the first onset of plastic, non-linear deformation in the F_n,ens_ and F_n,pos_ curves at x_pos_ = 2.8%, y_pos_ = 2.17, x_ens_ = 14.4% and y_ens_ = 1.16. The linear interpolation between these two injury threshold points (Eq 1) was then used to find the minor TSIC thresholds for the EHTM and TMM as the tendon strain at the intersection between the normalised EHTM and TMM tendon force curves with the linearly interpolated injury threshold line (Eq 4). This process is depicted in Fig 4. The equation describing the linearly interpolated threshold line is given in Eq 10. The corresponding minor TSIC threshold values of positional, TMM, EHTM and energy storage tendons, as well as all other curve parameters are listed in Table 1. The derived TSIC thresholds, together with the inherently scaling MSIC thresholds, were used for the strain injury assessments presented in the following parts of this work.

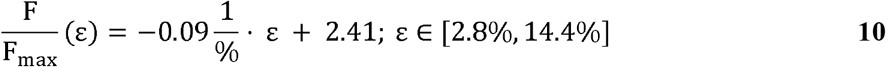

where F is the tendon force, F_max_ is the connected muscles maximum isometric force and ε is the tendon strain in percent.

**Fig 4.**
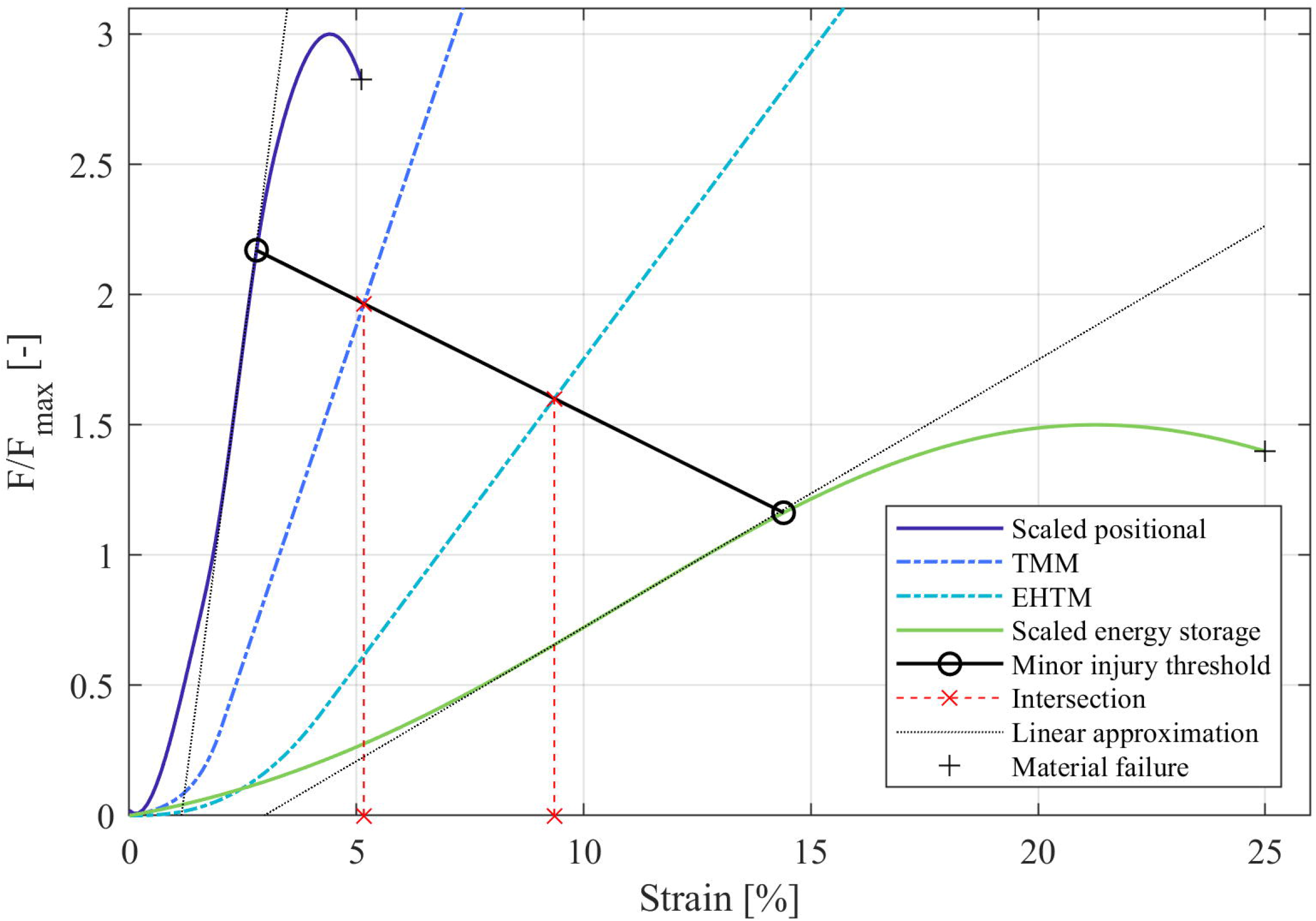
Normalised and scaled force-strain curves of positional and energy storage tendons compared to the normalised force-strain curves of EHTM and TMM tendons.

## 3 Results

### 3.1 Strain injury assessment during repositioning

The THUMS repositioning simulation terminated without errors of any kind, indicating that no severe mesh deformation occurred during the runtime. The strain injury assessment of the 18 EHTM muscles showed that both the left- and right-hand *musculus pronator teres* sustained minor muscle strain injury at t = 1.04 s (Fig 5, S-Fig 4 in S1 File). This straining of the muscle coincides with the widening of the elbow joint gap during repositioning, which increased from 6.30 mm at t = 0.0 s to 12.92 mm at t = 1.2 s (Fig 6, S-Fig 5 in S1 File). Comparing these joint gap values to experimental data from Lee et al. [78], where average elbow joint gaps of 3.3 ±0.4 mm were measured, shows that the initial joint gap in the THUMS model is already unphysiologically wide. This modelling error is exacerbated by the repositioning, where the extension of the arms widened the joint gap by another 6.62 mm. Measurements of elbow joint gaps in healthy and dislocated elbows by Hopf et al. [79] revealed medial joint gap differences of merely 1.6 ±1.1 mm, indicating that the repositioning simulation effectively dislocated the elbow joint multiple times over. Through the injury assessment of muscles spanning the joint gap, we were able to identify this internal model deformation, which would be almost indetectable by conventional automated mesh quality assessment tools.

**Fig 5.**
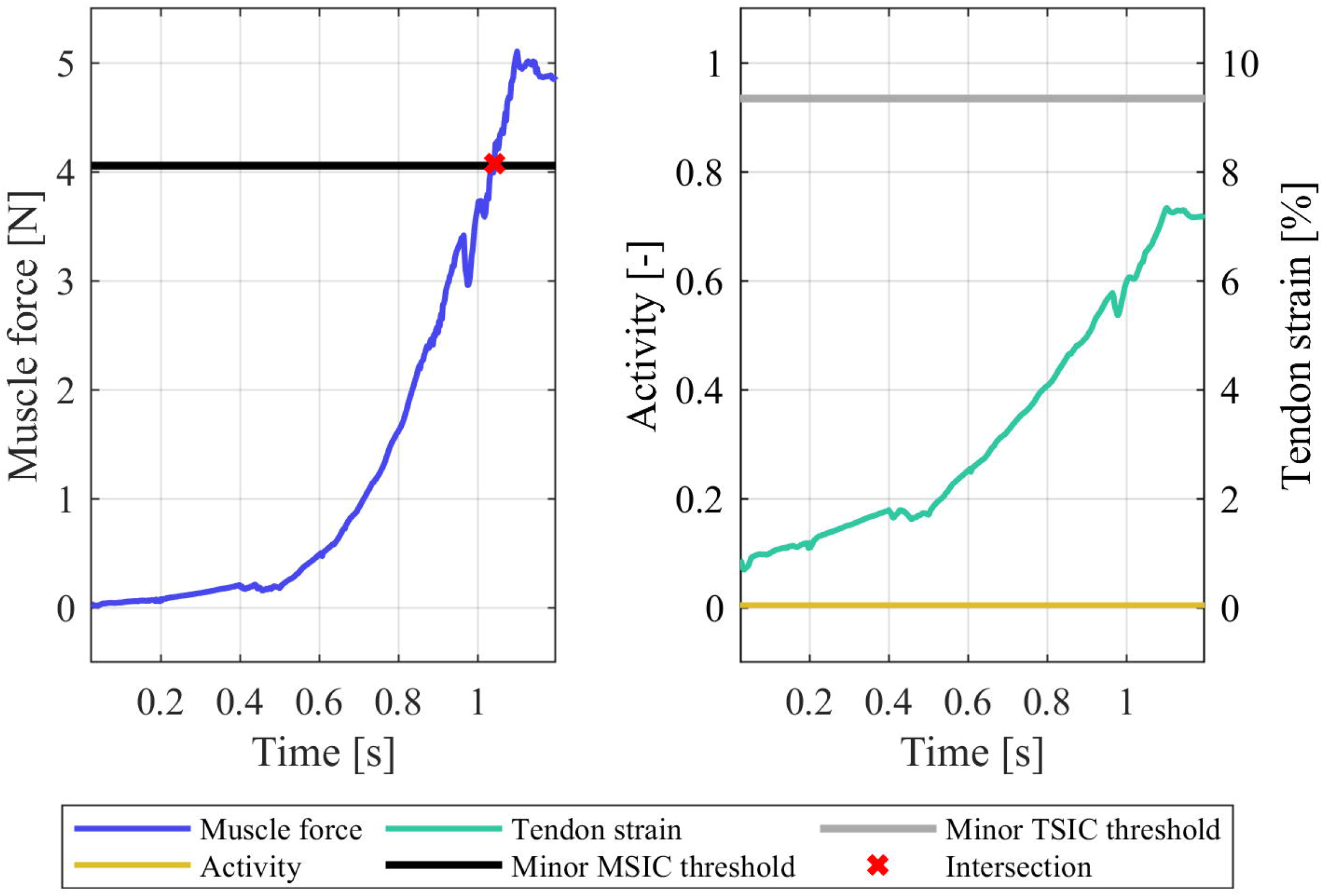
Strain injury assessment results of the left-hand *musculus pronator teres*.

**Fig 6.**
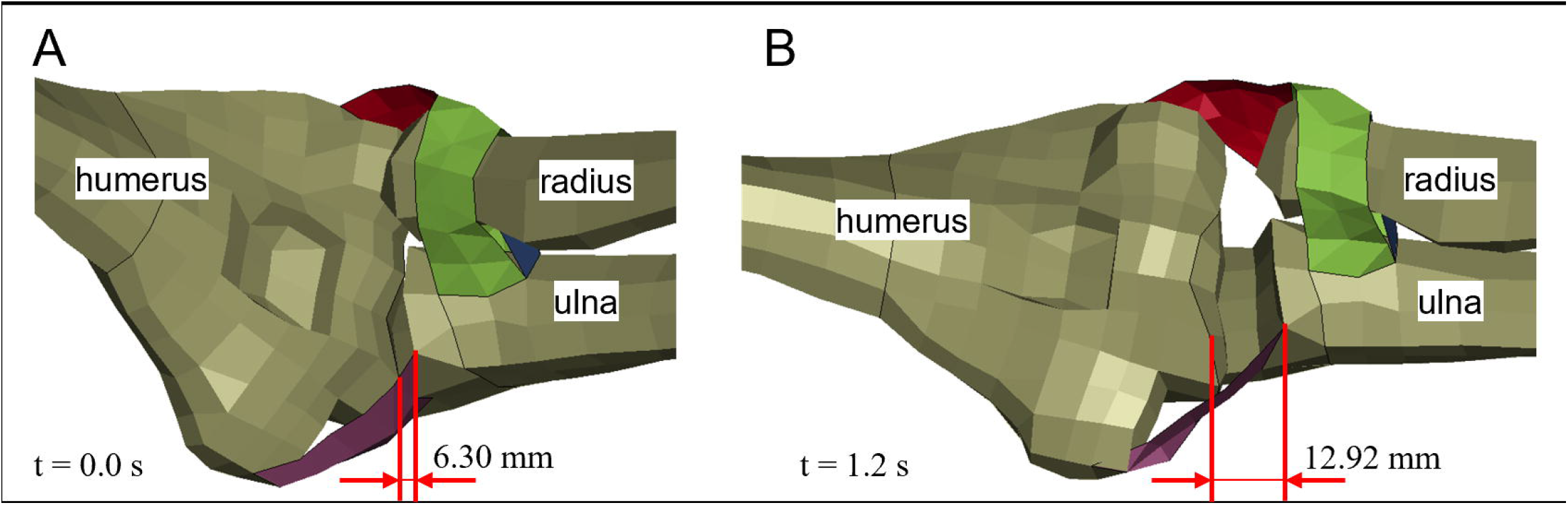
Joint gap in the left elbow joint of the THUMS version 5.03 occupant model. (A) Initial THUMS occupant position at t = 0.0 s ; (B) Repositioned THUMS with extended arms at t = 1.2 s.

### 3.2 Strain injury assessment under varying muscle material parameterisations

The strain injury assessment of the partial gait cycle simulations with the gait2354 model showed no injuries for the default model (Fig 7). However, in the case of the gait2354 model with modified *musculus gastrocnemius medialis* parametrisation, both minor muscle and tendon strain injuries occurred in the differently configured muscle (Fig 8). The shortening of the tendon slack length led to a considerable pre-strain in the tendon (3.84% at t = 0.0 s) which created injurious tensile forces in the muscle. The elongation of the MTU during the gait cycle then led to a further elongation of the tendon, crossing the minor TSIC threshold of 5.16% strain at t = 1.03 s and reaching a maximum strain of 5.45% at the termination time of t = 1.08 s. Even though the model kinematics are virtually indistinguishable between the default and modified gait2354 models (Fig 2, S-Fig 6 in S1 File), the proposed method for detecting erroneously parameterised muscles via strain injury assessment was able to pinpoint the modified *musculus gastrocnemius medialis* while simultaneously confirming the parameter validity of the default gait2354 model.

**Fig 7.**
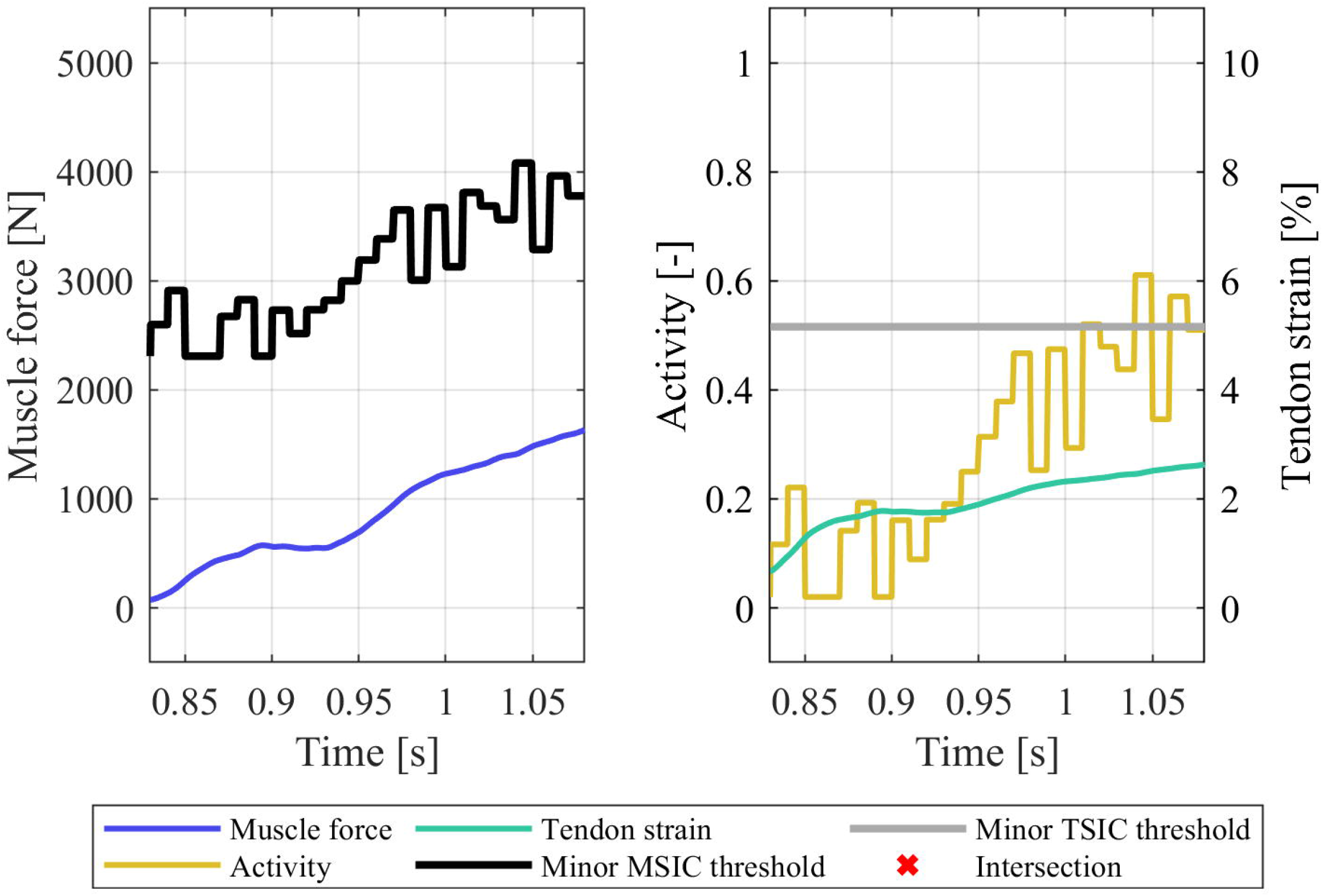
Strain injury assessment results of the right-hand *musculus gastrocnemius medialis* in the default gait2354 model.

**Fig 8.**
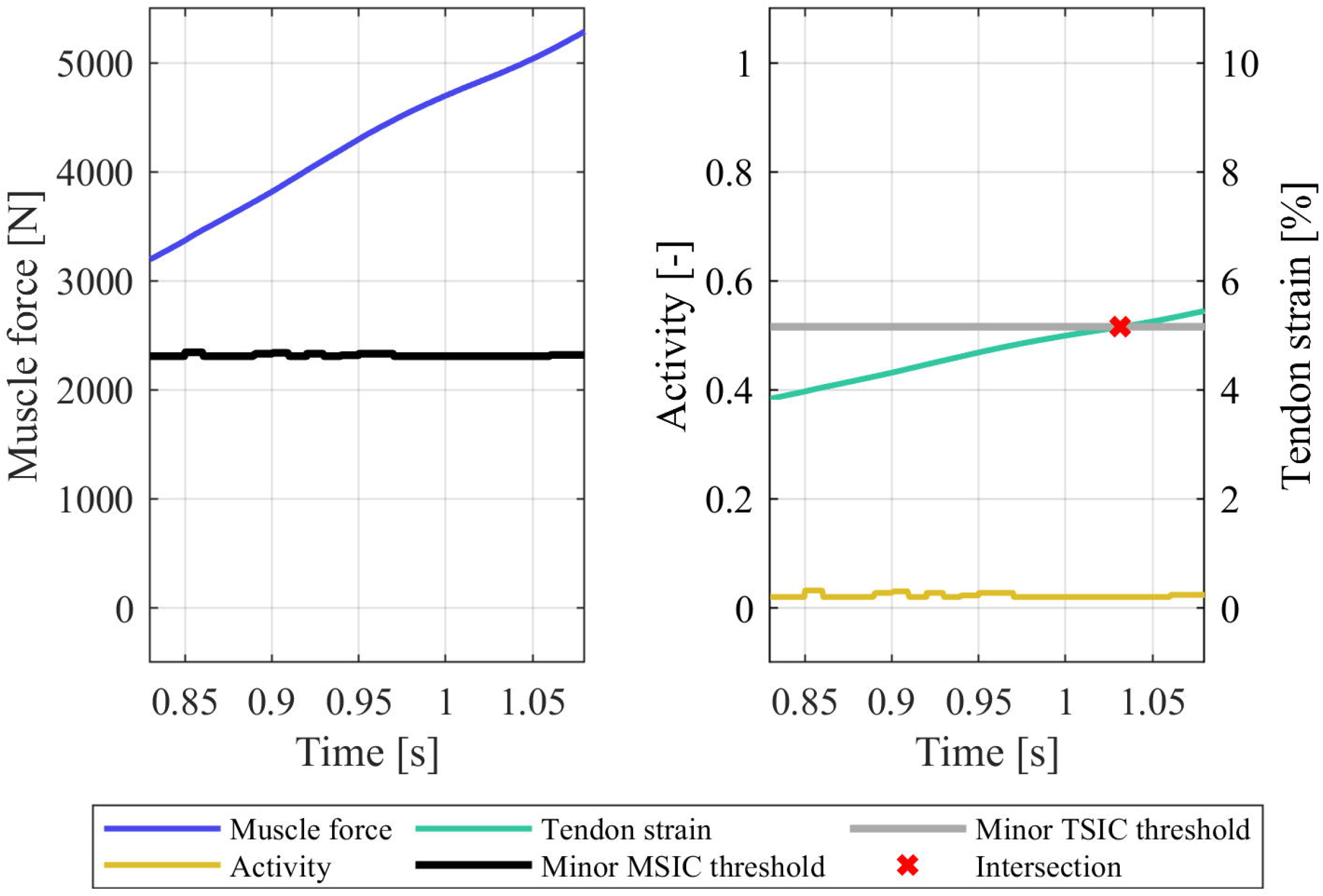
Strain injury assessment results of the right-hand *musculus gastrocnemius medialis* in the modified gait2354 model.

## 4 Discussion

In our work, the exemplary repositioning of the THUMS model was performed by applying external forces to the FE model to adjust its posture. While some might consider this method outdated [13], it remains representative of how FE models are repositioned to this day [80,81]. We therefore consider our choice of repositioning approach to be justified and relevant to the presented issues at hand. The force based MSIC thresholds are highly sensitive to quick strains and fast contraction velocities of the muscle, as dampening elements within the EHTM can produce high forces if severe length rate changes occur. Consequently, the repositioning speeds and model settling times were chosen such that peak material stresses were avoided. The measurements of the elbow joint gap during repositioning were done according to the methodology of Lee et al. [78] to ensure comparability with their results. As the joint gap in dislocated elbows was determined differently by Hopf et al. [79], we only compared the detected change in joint gap as opposed to absolute values.

For the gait cycle MB simulations in OpenSim, the CMC functionality was used to derive the control signals for the following forward dynamics simulations. This option was chosen, as other methods to find muscle stimulation signals, such as static optimisation [82], do not account for the deformation of passive mechanical structures of the muscle. Accordingly, no tendon strains can be derived from static optimisation, making it impossible to apply the chosen injury criteria. The duration of the simulated gait cycle was predetermined by the simulation setups provided with OpenSim [59,60]. While full gait cycle simulations or simulations with larger ranges of motion, like squats, would have shown a more complete picture of the MTU strains in a typical movement scenario, the authors prioritised reproducibility and thus opted for the use of openly available and well-documented example simulations instead. Additionally, the simulated partial gait cycle was sufficient to illustrate the systemic errors, such as large pre-strains of the MTU, which can occur if the corresponding Hill-type parameters are chosen poorly.

A possible point of uncertainty is the scaling of the force-strain curves used for determining tendon specific injury thresholds to ensure consistency between the TSIC and MSIC thresholds and to align the tendon deformation characteristics with the findings of previous works [75–77]. This retroactive scaling of experimental data was deemed necessary as no publication listing all required parameters for the transfer of a stress-strain curve to a normalised force-strain curve is known to the authors. A study by Knaus et al. [83] has shown that Achilles tendon CSA grows with the volume of the connected muscles and thus their maximum force production capability. Thus, a possible source of uncertainty could be that the CSA and F_max_ values we derived from different literature sources. However, any possible variation in tendon CSA could not reasonably explain the observed F_ens/pos_ to F_max_ ratios. For the calculated F_max,ens_ of 5500.3 N, a CSA_ens_ value of 124.93 mm^2^ would be required to achieve the desired max(F_ens_) to F_max_ ratio of 1.5. This is more than double the Achilles tendon CSAs measured by Kruse et al. (60.9 ± 8.2 mm^2^) [71] and Knaus et al. (64.8 ± 16.34 mm^2^) [83], which is beyond the magnitude of any measurement error or standard deviation. On the other hand, assuming a correctly set CSA of 60.9 mm^2^ and calculating the theoretical F_max_ based on the same target of max(F_ens_) = 1.5 F_max_ would indicate that the maximum force acting on the Achilles tendon could not exceed 2681.25 N, less than the F_max_ of just the connected *musculus soleus* (3585.9 N [74]).

Moreover, the material characteristics of tendons described in literature are partly contradictory, further complicating the task of identifying correct material parameters. For example, Achilles tendon ultimate tensile strengths ranging from 48 MPa [68] to 86 MPa [69] can be found, while experiments by Komi et al. [84] have shown that stresses of up to 111 MPa act on the tendon during sprinting. Similarly, Fukashiro et al. measured Achilles tendon forces of 3787 N during hopping [85]. Comparing this measurement to our originally calculated ultimate Achilles tendon strength of 4022 N (Eq 6) would indicate that hopping introduces a load of 94% ultimate strength in the Achilles tendon. Given these apparent contradictions and the wide spread of tendon material properties described in the literature [86], the authors feel justified in scaling the force-strain curves to reflect biomechanically sensible injury resistances. Still, these assumptions on our part are a limitation of the presented work. However, the exact TSIC threshold values derived from Fig 4 are not essential to the model assessment method described within this manuscript. If a more complete data set of tendon material parameters is know to any reader, new boundary curves for the tendon compliance corridor and TSIC thresholds can be defined using the generalised calculation steps outlined in Eq 1, Eq 2, Eq 3 and Eq 4.

In addition to providing a method for deriving minor TSIC thresholds, Fig 4 also enables us to qualitatively assess towards which extreme tendon archetype a model’s tendons conform most to. The TMM tendons are comparatively stiff (slope 0.52/% compared to energy storage tendon slope of 0.10/%) given that they supposedly represent tendons involved in locomotion, where the energy storage characteristics of tendons would assuredly become relevant. On the other hand, the EHTM tendons are rather compliant (slope 0.24/% compared to positional tendon slope of 1.33/%), which could be considered atypical for the upper arm region. We can thus conclude that future works with the TMM and EHTM muscle models should reassess their general choice of tendon material parameters to better reflect the tendons’ mechanical purpose in the human body.

The modelling and parametrisation issues highlighted in this paper are by no means unavoidable. The deformations of FE models during repositioning could be significantly reduced if relevant anatomical structures were modelled in their entirety. The THUMS version 5.03 elbow is missing the lateral ulnar collateral ligament, the accessory collateral ligament and any form of joint capsule [87]. Introducing these elements might improve the joint deformation behaviour. Additionally, ligaments could be artificially stiffened during the repositioning simulations to further reduce the risk of joint dislocation. Similarly, adhering to standard methods of Hill-type muscle parameter tuning [88] will almost assuredly alleviate the issue of high MTU pre-strains, if the full range of motion of the muscle is taken into account during the tuning process.

### 4.1 Conclusions

The results of this work show that the proposed method can quantify the internal deformation behaviour of musculoskeletal models and the validity of Hill-type muscle parameter choice via strain injury assessment. However, the MSIC and TSIC thresholds are only capable of acting as an upper bound for muscle forces and tendon strains. If a model’s musculature were to be set-up in such a way as to show slack muscles throughout entire ranges of motion, no injuries of any kind could be detected. Consequently, the proposed method would indicate a physiological model behaviour, even though no movement could be generated with it. Our method is thus not a wholistic assessment tool but rather a valuable addition to an arsenal of other model quality evaluation approaches. Considering that the necessary quantities to perform MSIC and TSIC assessments are in most cases automatically output during a simulation’s runtime, we strongly encourage the informed interpretation of muscle forces and tendon strains to gain a deeper insight into the behaviour of muscle driven human body models.

## Acknowledgements

The authors would like to thank Michael Günther for his help and his valuable input during the creation of this paper.

## Supporting information

### S1 File

