## Supplementary material for "Using muscle-tendon load limits to assess unphysiological musculoskeletal model deformation and Hill-type muscle parameter choice": S1 File

### *Supporting Information*

#### 1 Supplementary Tables

**S-Table 1.** Specific muscle parameters of all implemented hand muscles.

| **Muscle Name** | $F_{max}$[1] **[N]** | $l_{CE,opt}$ **[mm]** | $l_{SEE,0}$ **[mm]** | $\Delta F_{SEE,0}$ **[N]** |
|---|---|---|---|---|
| *Musculus brachialis* | 987.30 | 111.45 | 69.98 | 394.92 |
| *Musculus biceps brachii* long head | 15.36 | 130.16 | 194.31 | 6.14 |
| *Musculus biceps brachii* short head | 13.07 | 104.23 | 183.27 | 5.23 |
| *Musculus brachioradialis* | 27.03 | 217.34 | 48.56 | 10.81 |
| *Musculus triceps brachii* long head | 798.50 | 158.38 | 169.02 | 319.40 |
| *Musculus triceps brachii* lateral head | 624.30 | 151.98 | 130.65 | 249.72 |
| *Musculus triceps brachii* medial head | 624.30 | 129.76 | 103.58 | 249.72 |
| *Musculus anconeus* | 350.00 | 23.54 | 15.70 | 140.00 |
| *Musculus pronator teres* | 4.48 | 36.52 | 94.38 | 1.79 |

Abbreviations: $\Delta F_{SEE,0}$ = Force at the nonlinear–linear transition in $F_{SEE}(l_{SEE})$; $F_{max}$ = Maximum isometric force; $l_{CE,opt}$ = Optimal muscle fibre length; $l_{SEE,0}$ = Rest length of the serial elastic element;

**S-Table 2.** Generic muscle parameters of all implemented hand muscles.

| Variable | Unit | Value | Description | Reference |
|---|---|---|---|---|
| $q_0$ | [-] | 0.005 | Minimum value of muscle activity | [2] |
| $c$ | [mol/L] | 1.37e-4 | Hatze constant $c$ | [3] |
| η | [L/mol] | 5.27e4 | Hatze constant η | [3] |
| $k$ | [-] | 2.9 | Hatze constant $k$ | [3] |
| $m$ | [1/s] | 11.3 | Hatze constant $m$ | [3] |
| $\Delta W_{des}$ | [-] | 0.45 | Width of $F_{isom}(l_{CE})$ on descending limb | [4] |
| $\nu_{CE,des}$ | [-] | 1.5 | Exponent of $F_{isom}(l_{CE})$ on descending limb | [5] |
| $\Delta W_{asc}$ | [-] | 0.45 | Width of $F_{isom}(l_{CE})$ on ascending limb | [4] |
| $\nu_{CE,asc}$ | [-] | 3 | Exponent of $F_{isom}(l_{CE})$ on ascending limb | [5] |
| $A_{rel,0}$ | [-] | 0.2 | Maximum value of $A_{rel}$ | [2] |
| $B_{rel,0}$ | [1/s] | 2.0 | Maximum value of $B_{rel}$ | [2] |
| $S_{ecc}$ | [-] | 2.0 | Step in inclination of $F_{CE}(\dot{l}_{CE} = 0)$ between eccentric and concentric force-velocity relations | [6] |
| $F_{ecc}$ | [-] | 1.5 | Coordinate of pole in $l_{CE}(F_{CE})$ normalised to $F_{max} q F_{isom}(l_{CE})$ for $l_{CE} > 0$ | [6] |
| $L_{PEE,0}$ | [-] | 0.95 | Rest length of the PEE normalised to $l_{CE,opt}$ | [2] |
| $\nu_{PEE}$ | [-] | 2.5 | Exponent of $F_{PEE}(l_{CE})$ | [5] |
| $F_{PEE}$ | [-] | 2.0 | Force of PEE if $l_{CE}$ is stretched to $\Delta W_{des}$ | [5] |
| $\Delta U_{SEE,nll}$ | [-] | 0.0425 | Relative stretch at non-linear-linear transition in $F_{SEE}(l_{SEE})$ | [5] |
| $\Delta U_{SEE,l}$ | [-] | 0.017 | Relative stretch in linear part for force increase $\Delta F_{SEE,0}$ | [5] |
| $D_{SDE}$ | [-] | 0.3 | Dimensionless factor to scale $d_{SE,max}$ | [5] |
| $R_{SDE}$ | [-] | 0.01 | minimum value of $d_{SE}$ normalised to $d_{SE,max}$ | [5] |

#### 2 Supplementary Figures

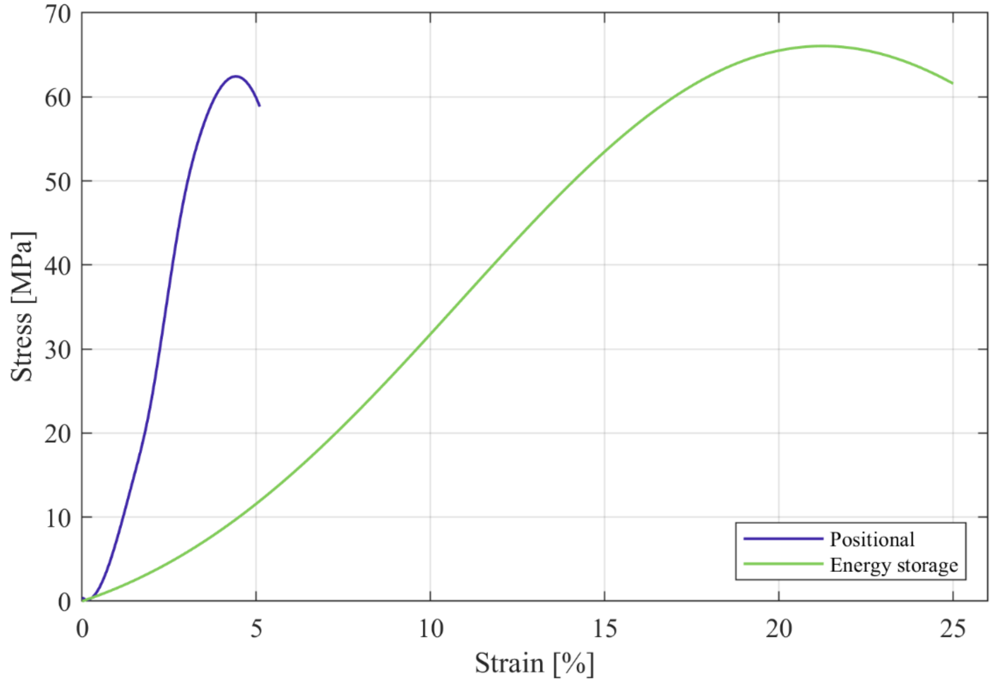


**S-Fig 1.** Stress-strain curves of positional and energy storage tendons from literature [7,8].

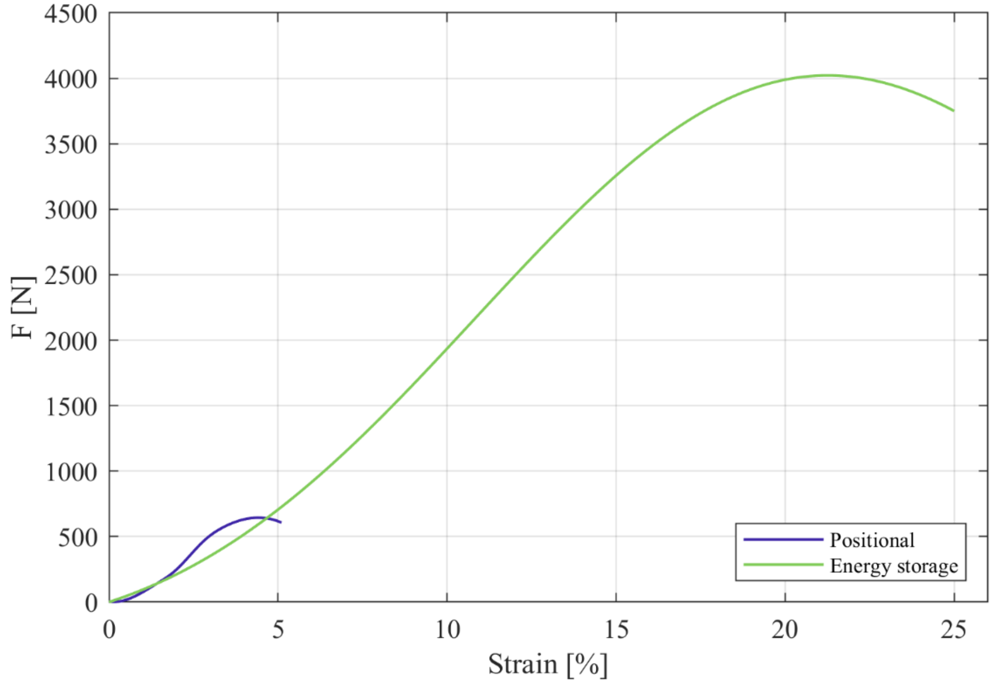


**S-Fig 2.** Force-strain curves of positional and energy storage tendons.

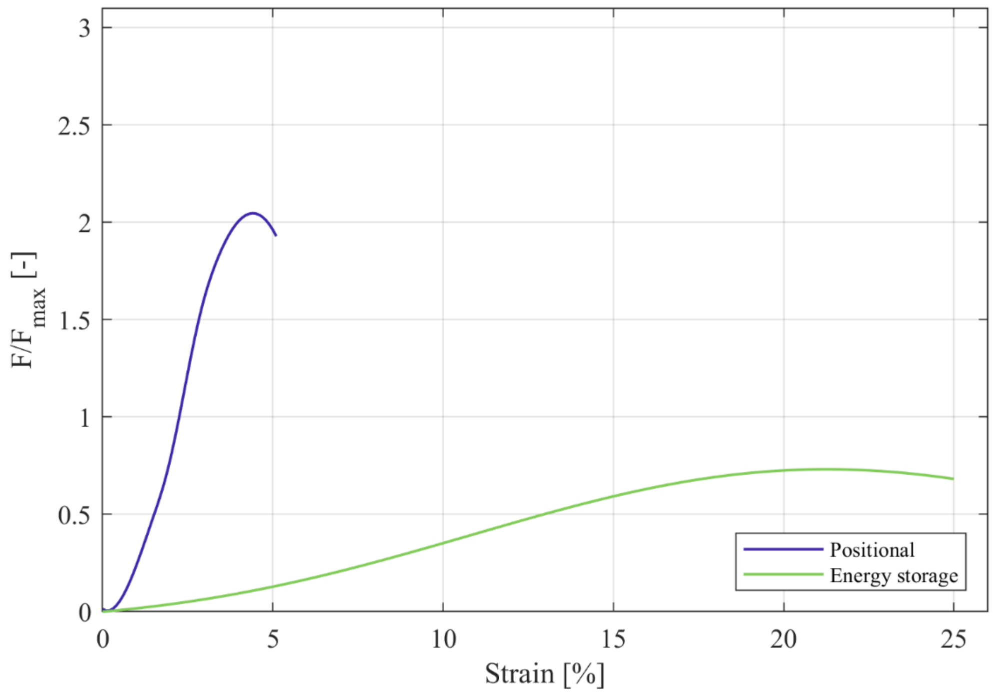


**S-Fig 3.** Normalised force-strain curves of positional and energy storage tendons.

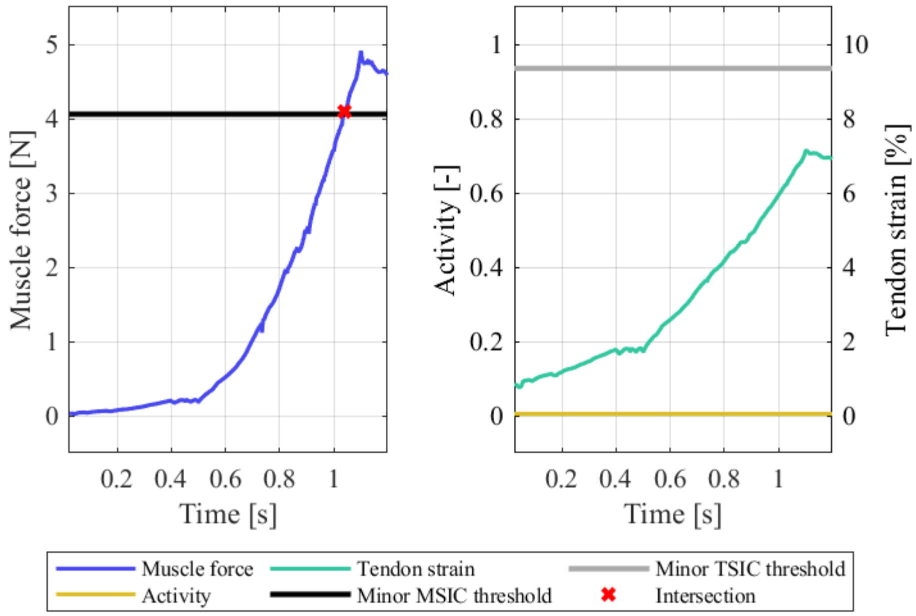


**S-Fig 4.** Strain injury assessment results of the right-hand *musculus pronator teres.*

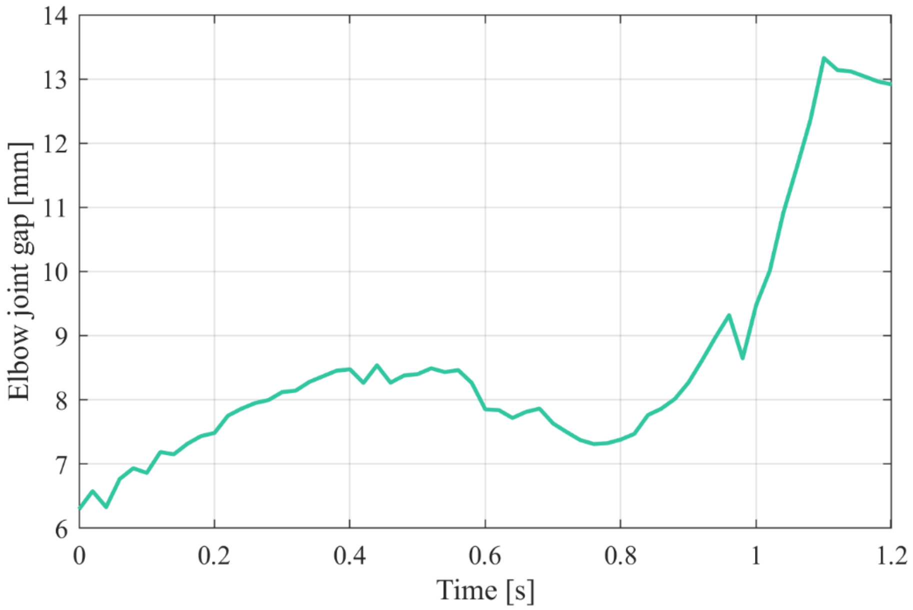


**S-Fig 5.** Joint gap in the left-hand elbow joint during the THUMS repositioning simulation.

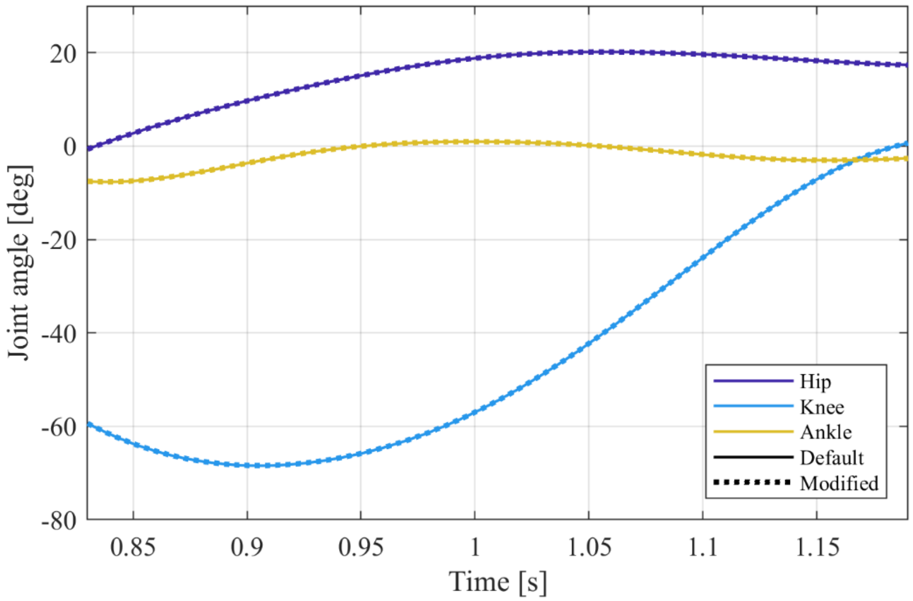


**S-Fig 6.** Comparison of the hip, knee, and ankle joint angles in sagittal plane during the partial gait cycle simulation using the default and modified gait2354 models.
